# Abolishing the prelamin A ZMPSTE24 cleavage site leads to progeroid phenotypes with near-normal longevity in mice

**DOI:** 10.1101/2021.11.11.465186

**Authors:** Yuexia Wang, Khurts Shiladardi, Trunee Hsu, Kamsi O. Odinammadu, Takamitsu Maruyama, Wei Wu, Chyuan-Sheng Lin, Christopher B. Damoci, Eric D. Spear, Ji-Yeon Shin, Wei Hsu, Susan Michaelis, Howard J. Worman

## Abstract

Prelamin A is a farnesylated precursor of lamin A, a nuclear lamina protein. Accumulation of the farnesylated prelamin A variant progerin, with an internal deletion including its processing site, causes Hutchinson-Gilford progeria syndrome. Loss of function mutations in *ZMPSTE24*, which encodes the prelamin A processing enzyme, lead to accumulation of full-length farnesylated prelamin A and cause related progeroid disorders. Some data suggest that prelamin A also accumulates with physiological aging. *Zmpste24*^-/-^ mice die young, at ~20 weeks. Because ZMPSTE24 has functions in addition to prelamin A processing, we generated a mouse model to examine effects solely due to the presence of permanently farnesylated prelamin A. These mice have an L648R amino acid substitution in prelamin A that blocks ZMPSTE24-catalyzed processing to lamin A. The *Lmna*^L648R/L648R^ mice express only prelamin and no mature protein. Notably, nearly all survive to 65-70 weeks, with approximately 40% of male and 75% of female *Lmna*^L648R/L648R^ mice having near-normal lifespans of 90 weeks (almost 2 years). Starting at ~10 weeks of age, Lmna^L648R/L648R^ mice of both sexes have lower body masses and body fat than controls. By ~20-30 weeks of age, they exhibit detectable cranial, mandibular and dental defects similar to those observed in *Zmpste24*^-/-^ mice, and have decreased vertebral bone density compared to age- and sex-matched controls. Cultured embryonic fibroblasts from *Lmna*^L648R/L648R^ mice have aberrant nuclear morphology that is reversible by treatment with a protein farnesyltransferase inhibitor. These novel mice provide a robust model to study the effects of farnesylated prelamin A during physiological aging.

## Introduction

The lamin A/C gene (*LMNA*) encodes the splice variants lamin A and lamin C, which are intermediate filament building blocks of the nuclear lamina that differ in their carboxyl-terminal domains. Prelamin A, but not lamin C, has a carboxyl-terminal cysteine-aliphatic-aliphatic-any amino acid (CAAX) motif that initiates a series of posttranslational processing reactions to generate mature lamin A. In addition to farnesylation, cleavage of -AAX, and carboxymethylation (CAAX processing), prelamin A undergoes a final cleavage reaction, uniquely catalyzed by the zinc metalloprotease ZMPSTE24. This removes the last 15 amino acids of prelamin A, including its farnesylated cysteine, resulting in the production of mature, unfarnesylated lamin A. Defects in the posttranslational processing of prelamin to lamin A cause progeroid disorders (Fig. 1*A*) (1–3).

**Fig. 1.**
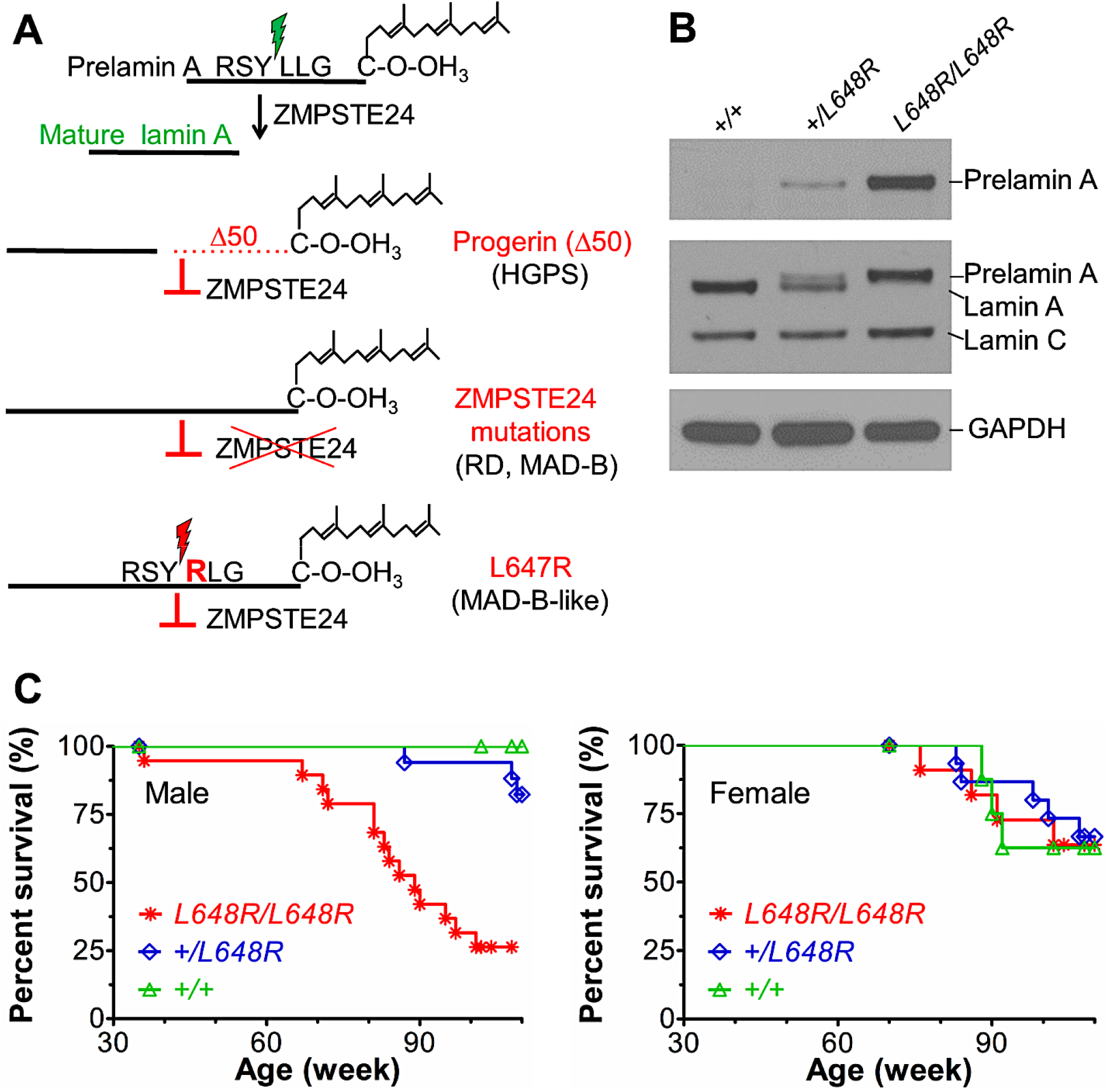
Survival of mice with a *Lmna* L648R mutation corresponding to the human mutation *LMNA* L647R that encodes an uncleavable variant of prelamin A. (*A*) Prelamin A is normally processed to lamin A after proteolytic cleavage catalyzed by the zinc metalloprotease ZMPSTE24 between tyrosine (Y) and leucine (L) 647 (648 in mouse), removing the carboxyl-terminal farnesylated cysteine (C). HGPS-causing *LMNA* mutations generate a prelamin A variant with an internal deletion of 50 amino acids (Δ50) called progerin, which lacks the ZMPSTE24 cleavage site and retains a farnesylated carboxyl-terminal cysteine. *ZMPSTE24* loss-of-function mutations cause restrictive dermopathy (RD) or MAD-B, in which unprocessed farnesylated prelamin A accumulates. *LMNA* mutation causing a MAD-B-like disorder generates a leucine (L) to arginine (R) substitution at residue 647 of prelamin A blocks ZMPSTE24 processing, leading to expression of a farnesylated variant with only a single amino acid difference. (*B*) Immunoblots of protein extracts from livers of *Lmna*^+/+^ (*+/+*), *Lmna*^+/L648R^ (*+/L648R*) and *Lmna*^L648R/L648R^ (*L648R/L648R*) mice. Blots were probed with an antibody specific for prelamin A (top), an anti-lamin A/C antibody that recognized prelamin A, lamin A and lamin C (middle) or anti-GAPDH antibody as loading control (bottom). (*C*) Survival curves for male *L648R/L648R* (*N* = 19), *+/L648R* (*N* = 17) and *+/+* (*N* = 10) mice and female *L648R/L648R* (*N* = 11), *+/L648R* (*N* = 15) and +/+ (*N* = 8) mice.

Hutchinson-Gilford progeria syndrome (HGPS) results from a splicing mutation in *LMNA* that generates a variant with a 50-amino acid internal deletion (Δ50) called progerin, which retains its CAAX motif but lacks the ZMPSTE24 cleavage site, and thus progerin’s carboxyl-terminal cysteine remains permanently farnesylated (4, 5). Children with HGPS manifest numerous premature aging symptoms, including failure to thrive, bone loss, and early-onset severe atherosclerosis leading to death typically in the second decade. Progeroid disorders also result from mutations in *ZMPSTE24* that lead to accumulation of full-length farnesylated prelamin A. Mandibuloacral dysplasia type B (MAD-B) is generally less severe than HGPS and caused by partial loss-of-function mutations in *ZMPSTE24*, with disease severity correlating with residual proteolytic activity (6, 7). Restrictive dermopathy, caused by complete loss-of-function mutations in *ZMPSTE24*, is neonatal lethal (8).

Genetically modified mouse models provide valuable tools to study these human progeroid disorders and may have the potential to illuminate the role of permanently farnesylated prelamin A in physiologic aging. *Zmpste24*^-/-^ mice accumulate farnesylated prelamin A and develop progeroid phenotypes, including severe growth retardation, craniofacial abnormalities, spontaneous bone fractures, and have a median survival of only approximately 20 weeks (9, 10). In *Zmpste24*^-/-^ mice, disease severity is ameliorated by genetic reduction of prelamin A or pharmacological treatment to block protein farnesylation (11–13). These findings, along with those showing a correlation between disease severity and residual enzyme activity in humans with *ZMPSTE24* mutations, suggest a “dose response” between the amount of farnesylated prelamin A and the degree of pathology. However, ZMPSTE24 has at least one other function besides prelamin A processing, namely clearing clogged translocons (14, 15). Hence, some of the organismal pathology caused by ZMPSTE24 deficiency may result from defects in protein translocation into the endoplasmic reticulum (ER). ZMPSTE24 may also have additional functions, inferred from its contribution to viral defense in mice (16, 17) and from genetic studies in yeast that have implicated it in ER protein quality control, membrane stress, and establishing membrane protein topology (18–22).

Substitution of a hydrophobic residue with an arginine at the ZMPSTE24 cleavage site in prelamin A (V637R in chickens and L647R in humans) blocks its proteolysis (23, 24), resulting in the accumulation of permanently farnesylated prelamin A. We have previously reported a patient with a heterozygous *LMNA* mutation generating the L647R amino acid substitution in prelamin A. This patient had a relatively mild progeroid disorder with clinical features similar to those of MAD-B (25). We therefore generated and characterized here mice with the corresponding L648R amino acid substitution in prelamin A to determine if *Lmna*^L648R/L648R^ mice have the same or different phenotype and disease severity as *Zmpste24*^-/-^ mice.

## Results

### Generation and survival of *Lmna* L648R mice

Normally, farnesylated prelamin A is a transient species, essentially undetectable in cells because of its efficient conversion to mature lamin A. However, *LMNA* mutations causing HGPS, *ZMPSTE24* mutations causing RD and MAD-B, and the *LMNA* L647R mutation causing a MAD-B-like disorder, lead to accumulation of farnesylated prelamin A or variants (Fig. 1*A*). Whereas several mouse lines with HGPS-causing and *Zmpste24* loss-of-function mutations have been generated and characterized (9, 10, 26–31), a mouse with a single amino acid substitution in prelamin A that causes a MAD-B-like disease in humans does not exist. Such a mouse could provide valuable information on the impact of prelamin A on promoting aging phenotypes, absent the complications in *Zmpste24-/-* mice, where altered processing of substrates in addition to prelamin A or other disrupted pathways may contribute to the phenotypes.

We therefore used a recombination-mediated genetic engineering strategy to generate mice carrying a *Lmna*^L648R^ allele (*SI Appendix*, Fig. S1*A*). It should be noted that L648R in mice is equivalent to L647R in humans. Founder mice carried one mutant and one wild type allele (*SI Appendix*, Fig. S1*B*). We crossed founder male mice to female *Lmna*^+/+^ C57BL/6J mice and intercrossed the heterozygous male and female offspring to generate heterozygous *Lmna*^+/L648R^ and homozygous *Lmna*^L648R/L648R^ mice. Male and female *Lmna*^L648R/L648R^ mice were able to breed and female mice were able to become pregnant up to approximately 24 weeks of age.

As expected, *Lmna*^+/+^ mice expressed mature lamin A and lamin C, *Lmna*^+/L648R^ mice expressed prelamin A, lamin A and lamin C and *Lmna*^L648R/L648R^ mice expressed prelamin A and lamin C, at roughly the same levels, but no mature lamin A (Fig. 1*B* and *SI Appendix*, Fig. S1*C*). For analysis of survival and growth as well as all subsequent experiments, we used mice that were crossed several generations onto a >90% pure C57BL/6J background. The homozygous *Lmna*^L648R/L648R^ mice had unexpectedly long lifespans, with nearly all living 65-70 weeks and ~40% of male and ~75% of female *Lmna*^L648R/L648R^ alive at 90 weeks (almost 2 years) of age (Fig. 1*C*).

### Growth and metabolic features of *Lmna* L648R mice

Heterozygous and homozygous *Lmna* L648R mice were grossly indistinguishable from their wild type littermates early in life. Starting at approximately 10 weeks of age, however, *Lmna*^L648R/L648R^ mice had decreased body masses and this difference became accentuated over time (Fig. 2*A*). Given the normal growth and survival of heterozygous *Lmna*^+/L648R^ mice, we performed more detailed phenotypic analyses only on the homozygous *Lmna*^L648R/L648R^ mice compared to *Lmna*^+/+^ wild type mice. Similarly, many other heterozygous *Lmna* mutant mice, including some heterozygous for the HGPS mutation, have modest, minimal, or in many cases no abnormal phenotypes (32, 33). Consistent with their decreased body mass compared to wild type mice with aging, male and female *Lmna*^L648R/L648R^ mice were visibly smaller than *Lmna*^+/+^ animals of the same age at 56 and 74 weeks (Fig. 2*B*).

**Fig. 2.**
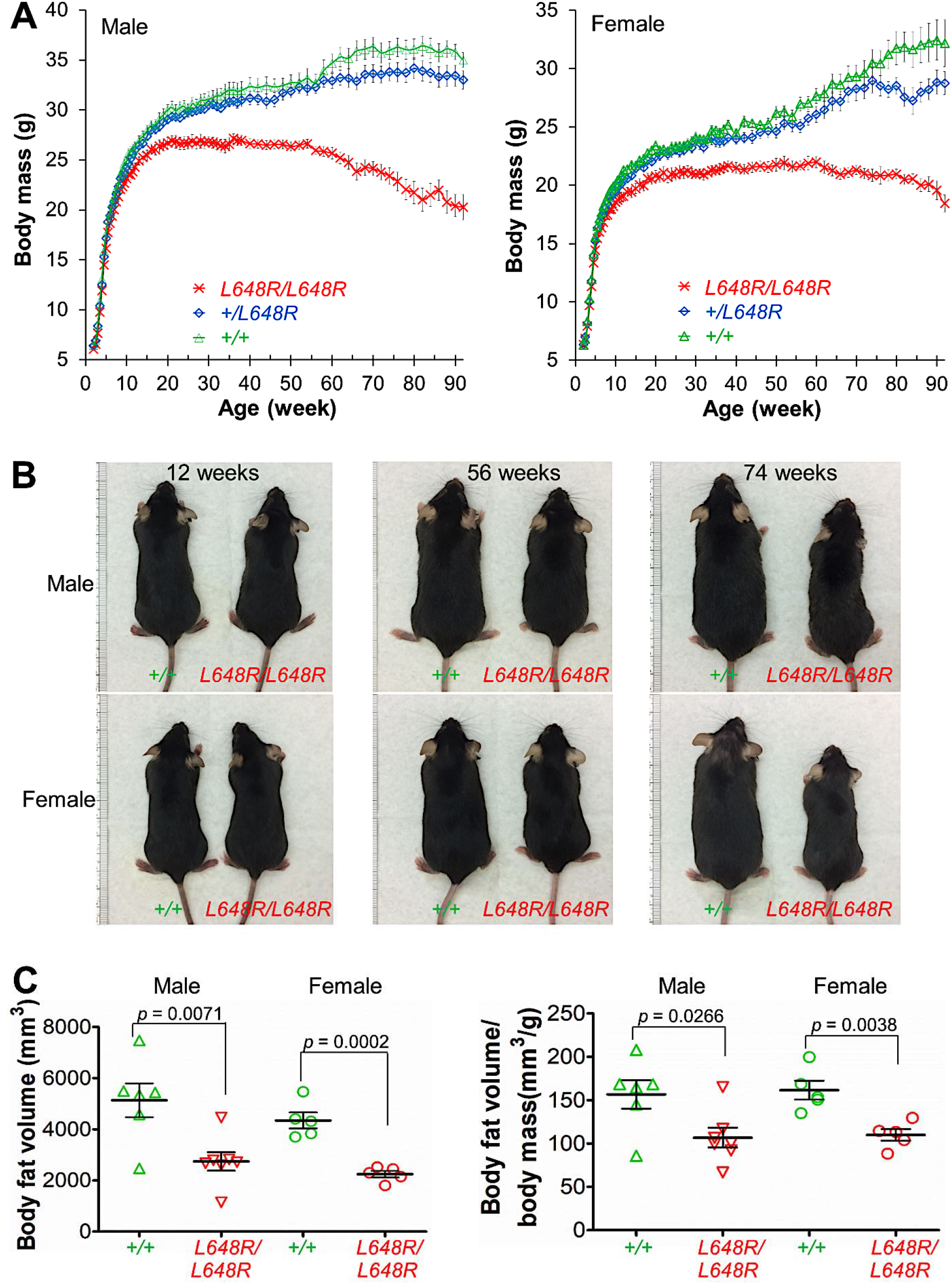
Growth of *Lmna*^L648R/L648R^ mice. (*A*) Body mass versus age of male *Lmna*^L648R/L648R^ (*L648R/L648R*) (*N* = 20), *Lmna*^+/L648R^ (*+/L648R*) (*N* = 17) and *Lmna^+/+^* (*+/+*) (*N* = 13) mice and female *L648R/L648R* (*N* = 13), *+/L648R* (*N* = 15) and +/+ (*N* = 11) mice. (*B*) Photographs comparing the sizes of male and female *+/+* and *L648R/L648R* mice at the indicated ages. (*C*) Body fat volume (left) and body fat volume normalized to body mass (right) of 52-week-old male and female *+/+* and *L648R/L648R* mice. Each triangle or circle represents value for an individual animal; long horizontal bars represent mean and errors bars indicate SEM.

*Zmpste24*^-/-^ mice have severe lipodystrophy along with growth retardation (9, 10). We therefore examined body fat composition and various metabolic parameters in *Lmna*^L648R/L648R^ mice. Male and female *Lmna*^L648R/L648R^ mice both had significantly reduced body fat at 52 weeks of age, compared to wild type mice (Fig. 2*C*). However, fasting blood glucose concentration at 30 weeks was not significantly different between *Lmna*^L648R/L648R^ and wild type mice (*SI Appendix*, Fig. S2*A*) at 30 weeks. Likewise, glucose tolerance testing showed no difference between male *Lmna*^L648R/L648R^ and *Lmna*^+/+^ mice, but the female mutant mice had decreased glucose tolerance compared to wild type controls (*SI Appendix*, Fig. S2*B*). Female *Lmna*^L648R/L648R^ also had lower plasma insulin concentrations compared to sex-matched *Lmna*^+/+^ mice (*SI Appendix*, Fig. S2*C*). There were no significant differences in other routine blood biochemical parameters between *Lmna*^L648R/L648R^ and *Lmna*^+/+^ mice (*SI Appendix*, Table S1). Thus, while *Lmna*^L648R/L648R^ mice clearly have low body mass and body fat, these phenotypes are not associated with any major prominent serum metabolic abnormality, except possible mild insulin resistance in female mice.

### Cranial and mandibular defects in *Lmna*^L648R/L648R^ mice

*Zmpste24*^-/-^ mice have profound cranial and mandibular abnormalities of the zygomatic arch, mandible, and dentition (9, 10). The human patient with the *LMNA* L647R mutation has microcephaly, micrognathia and dental crowding (25). We therefore examined the skulls of *Lmna*^L648R/L648R^ mice by micro-computed tomography (CT). At 4 weeks of age, skulls of male and female *Lmna*^L648R/L648R^ mice were essentially indistinguishable from *Lmna*^+/+^ mice. The zygomatic arches developed normally with the proper formation of both zygomaticotemporal and zygomaticomaxillary sutures in *Lmna*^L648R/L648R^ mice (Fig. *3A*). When rescanned at 30 weeks of age, however, male and female *Lmna*^L648R/L648R^ mice exhibited wide open zygomaticotemporal sutures in one or both zygomatic arches (Fig. 3*B*). These defects were present in one or both zygomatic arches in the majority of male and female *Lmna*^L648R/L648R^ mice but not in any *Lmna*^+/+^ mice at 30 weeks of age (Fig. 3*C*). The normal zygomatic arches 4 weeks after birth and defects at older ages implies that these are degenerative but not developmental deformities.

**Fig. 3.**
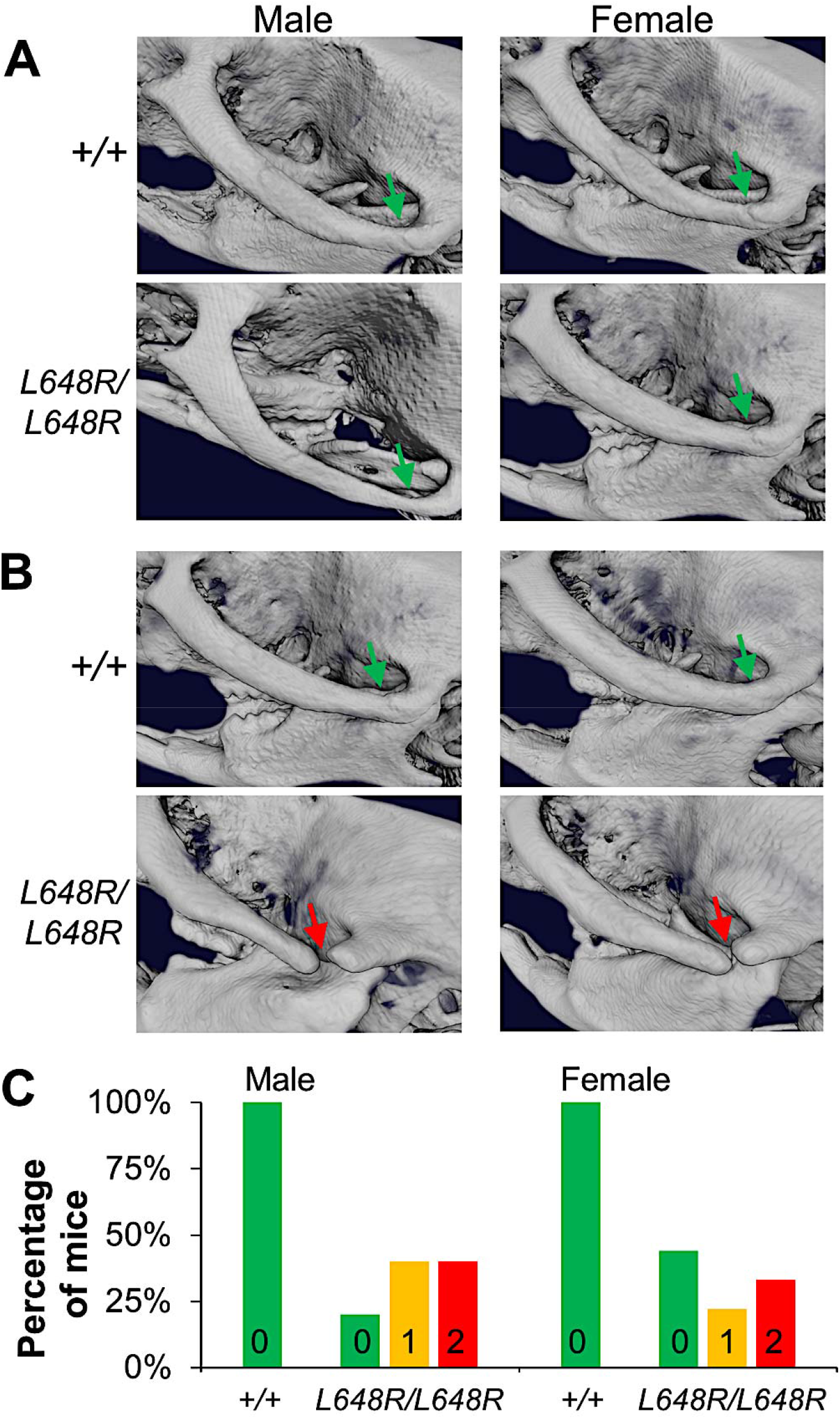
Degenerative deformities in zygomatic arches of *Lmna*^L648R/L648R^ mice. (*A*) Representative 3D rendering of the micro-CT images of living male and female *Lmna*^+/+^ (*+/+*) and *Lmna*^L648R/L648R^ (*L648R/L648R*) mice showing the formation of the zygomatic arches (green arrows) at 4 weeks. (*B*) Degenerative deformity (red arrows) in the zygomaticotemporal suture of male and female *L648R/L648R* mice compared to the proper maintenance (green arrows) in the *+/+* mice at 30 weeks. (*C*) Percentages of male *+/+* (*N* = 7), female *+/+* (*N* = 7), male *L648R/L648R* (*N* = 10) and female *L648R/L648R* (*N* = 9) mice with degenerative deformity in 0, 1 or 2 zygomatic arches at 30 weeks of age.

By 20 weeks of age, and more prominently at 30 weeks of age, micro-CT imaging and 3D-reconstruction analyses revealed profound mandibular degeneration in male and female *Lmna*^L648R/L648R^ mice (*SI Appendix*, Fig. S3). The 3D rendering of micro-CT scanned skull images after segmentation of mandible further examined the degenerative defects. The condylar, coronoid, and angular processes were severely deformed, resulting in decreased mandibular length, mandibular body length, and ramus height. (Fig. 4*A*). Mandible dimensions were significantly decreased in both male and female *Lmna*^L648R/L648R^ mice compared to age- and sex-matched *Lmna*^+/+^ controls (Fig. 4*B*). Dental malocclusion occurs in mice when the incisors overgrow because the mandibular and maxillary teeth are not normally aligned. Although oral examination showed the male and female *Lmna*^L648R/L648R^ had minimal to mild dental malocclusion at 30 weeks of age, this condition was quite evident at ages older than 70 weeks (Fig. 4*C*). This was confirmed by micro-CT scanning (*SI Appendix*, Fig. S4). By 65 weeks of age, virtually all male and 85% of female *Lmna*^L648R/L648R^ mice had dental malocclusion (Fig. 4*D*). This was likely attributed to the mandibular degeneration, leading to the improper alignment of the upper and lower jaws. *Lmna*^L648R/L648R^ mice clearly have mandibular and dental defects, which could compromise dietary intake and may contribute to decreased body mass observed with advancing age. The mandibular and dental abnormalities in *Lmna*^L648R/L648R^ mice resemble those of *Zmpste24*^-/-^ mice and those with HGPS mutations but appear less severe and are not prominent until later in life.

**Fig. 4.**
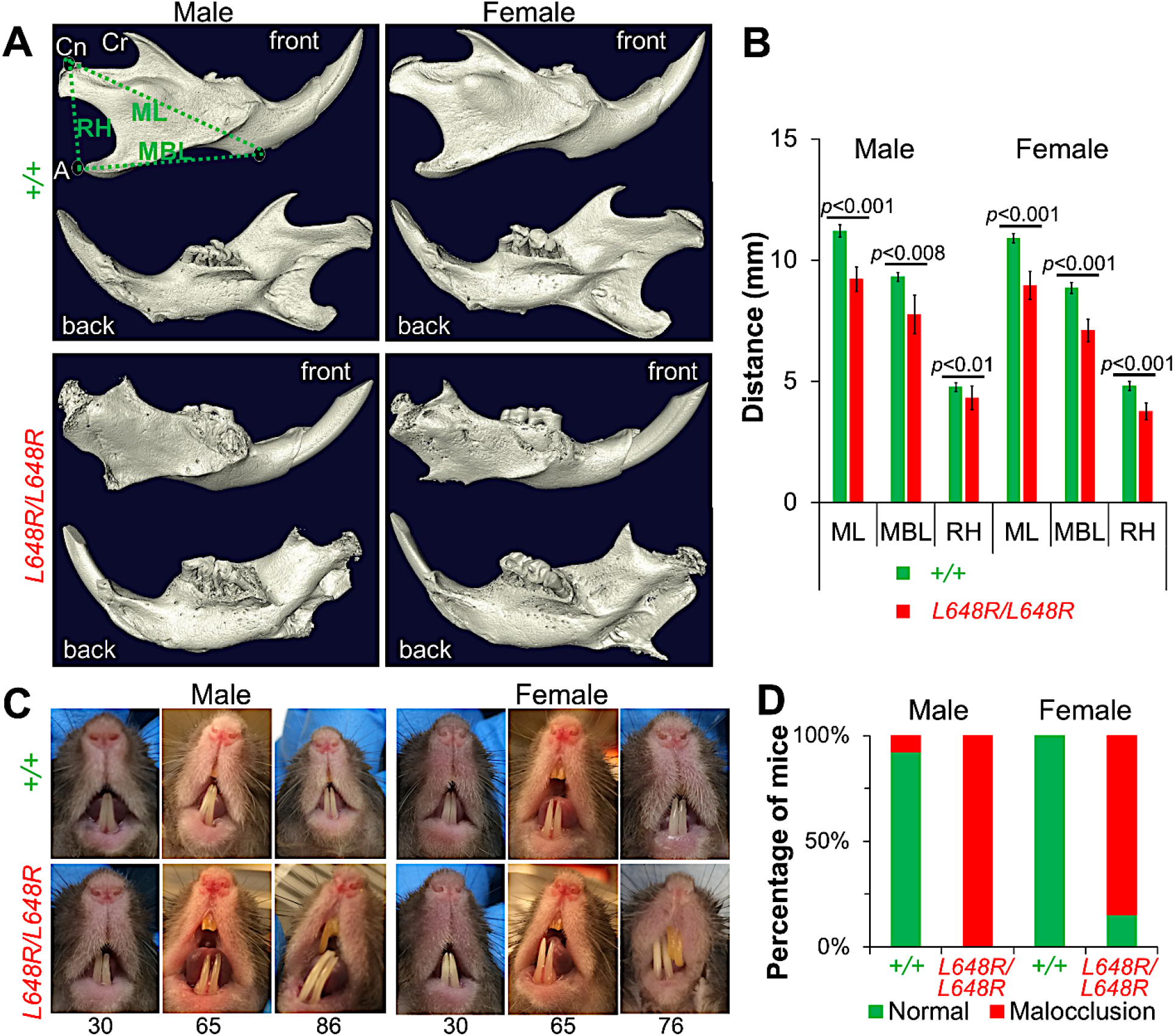
Mandibular defects in *Lmna*^L648R/L648R^ mice. (*A*) Representative 3D renderings of the segmented micro-CT-scanned images showing mandibles of male and female *Lmna*^+/+^ (*+/+*) and *Lmna*^L648R/L648R^ (*L648R/L648R*) mice. Cr: coronoid process; Cn: condylar process; A: angular process; ML: mandibular length; MBL: mandibular body length; RH: ramus height. (*B*) Comparisons of ML, MBL, and RH between male *+/+* (*N* = 4) and *L648R/L648R* (*N* = 4) mice and between female *+/+* (*N* = 4) and *L648R/L648R* (*N* = 4) mice. (*C*) Representative photographs of teeth of male and female *+/+* and *L648R/L648R* mice at the ages in weeks indicated. Dental malocclusion is minimal to mild at 30 weeks of age and more severe at older ages. (D) Percentages of male *+/+* (N = 13), female *+/+* (N = 10), male *L648R/L648R* (N = 18) and female *L648R/L648R* (N = 13) mice with malocclusion at ≥ 65 weeks of age.

### Decreased vertebral bone density and long bone defects in*Lmna*^L648R/L648R^ mice

*Zmpste24*^-/-^ mice suffer from osteoporosis and bone fractures (9, 10, 12, 34). We therefore used micro-CT scanned images to analyze vertebral bone density in *Lmna*^+/+^ and *Lmna*^L648R/L648R^ mice. From the 2D images, we first segmented the vertebra L5 by selecting the trabecular bone area. Next, we generated the 3D rendered image and computed statistical information of the 3D model to obtain bone and non-bone volumes (Fig. 5*A*). At 30 weeks of age, both male and female *Lmna*^L648R/L648R^ mice had significantly reduced vertebral bone density compared to sex-matched *Lmna*^+/+^ controls (Fig. 5*B*). Tibias of both male and female *Lmna*^L648R/L648R^ mice were thinner and had a more irregular surface than those of *Lmna*^+/+^ mice (Fig. 5*C*). These bone density abnormalities are similar to, but apparently less severe and of later onset, than those reported in *Zmpste24*^-/-^ mice.

**Fig. 5.**
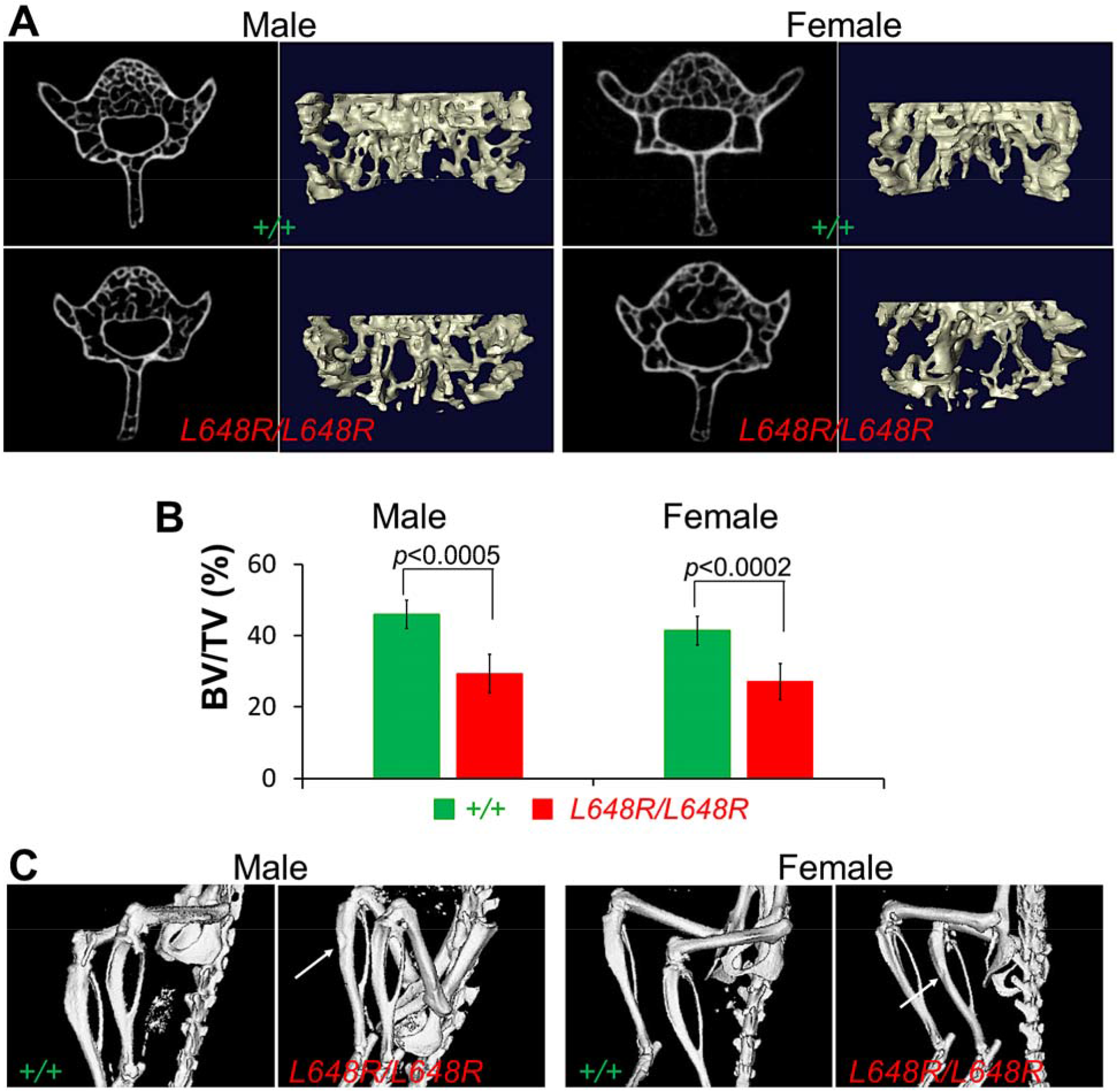
Decreased vertebral bone density and tibial defects in *Lmna*^L648R/L648R^ mice. (*A*) Representative micro-CT-scanned transverse sections and 3D reconstructions of the L5 vertebrae from male and female *Lmna*^+/+^ (*+/+*) and *Lmna*^L648R/L648R^ (*L648R/L648R*) mice. (*B*) Comparison of vertebra L5 bone density (bone volume/total volume; BV/TV %) between male *+/+* (*N* = 5) and *L648R/L648R* (*N* = 5) mice and between female *+/+* (*N* = 7) and *L648R/L648R* (*N* = 6) mice. (*C*) Micro-CT-generated representative images of hind legs of living male and female *+/+* and *L648R/L648R* mice. Arrows indicate thinner and more irregular surfaces of tibias of *L648R/L648R* mice.

### Farnesylation-dependent abnormal nuclear morphology in *Lmna*^L648R/L648R^ mouse embryonic fibroblasts (MEFs)

We isolated fibroblasts from *Lmna*^L648R/L648R^ and *Lmna*^+/+^ mouse embryos. Immunoblotting confirmed that *Lmna*^L648R/L648R^ MEFs produce only prelamin A and no mature lamin A, whereas the reverse is the case for *Lmna*^+/+^ MEFs. (Fig. 6*A*). Immunostaining similarly reveals that prelamin A accumulates in *Lmna*^L648R/L648R^ MEFs but not in *Lmna*^+/+^ MEFs (Fig. 6*B*). A high percentage of misshapen nuclei are apparent in mutant cells and this trend is amplified with passage number (Fig. 6*C*). Treatment of *Lmna*^L648R/L648R^ MEFs with the protein farnesyltransferase inhibitor (FTI) lonafarnib corrected the abnormal nuclear morphology (Fig. 6*D*). The percentage of mutant MEFs with abnormal nuclear morphology was significantly less in cultures treated with FTI compared to vehicle only (Fig. 6*E*). These results strongly suggest that accumulation of the farnesylated form of prelamin A is responsible for abnormal nuclear morphology. Given the correlations between FTI-induced reversal of abnormal nuclear morphology in MEFs from *Zmpste24*^-/-^ and HGPS mutant mice and the beneficial effects of these drugs on the corresponding whole animals (11, 31, 35–37) the farnesylated form of prelamin A also likely is responsible for the abnormal phenotypes in *Lmna*^L648R/L648R^ mice.

**Fig. 6.**
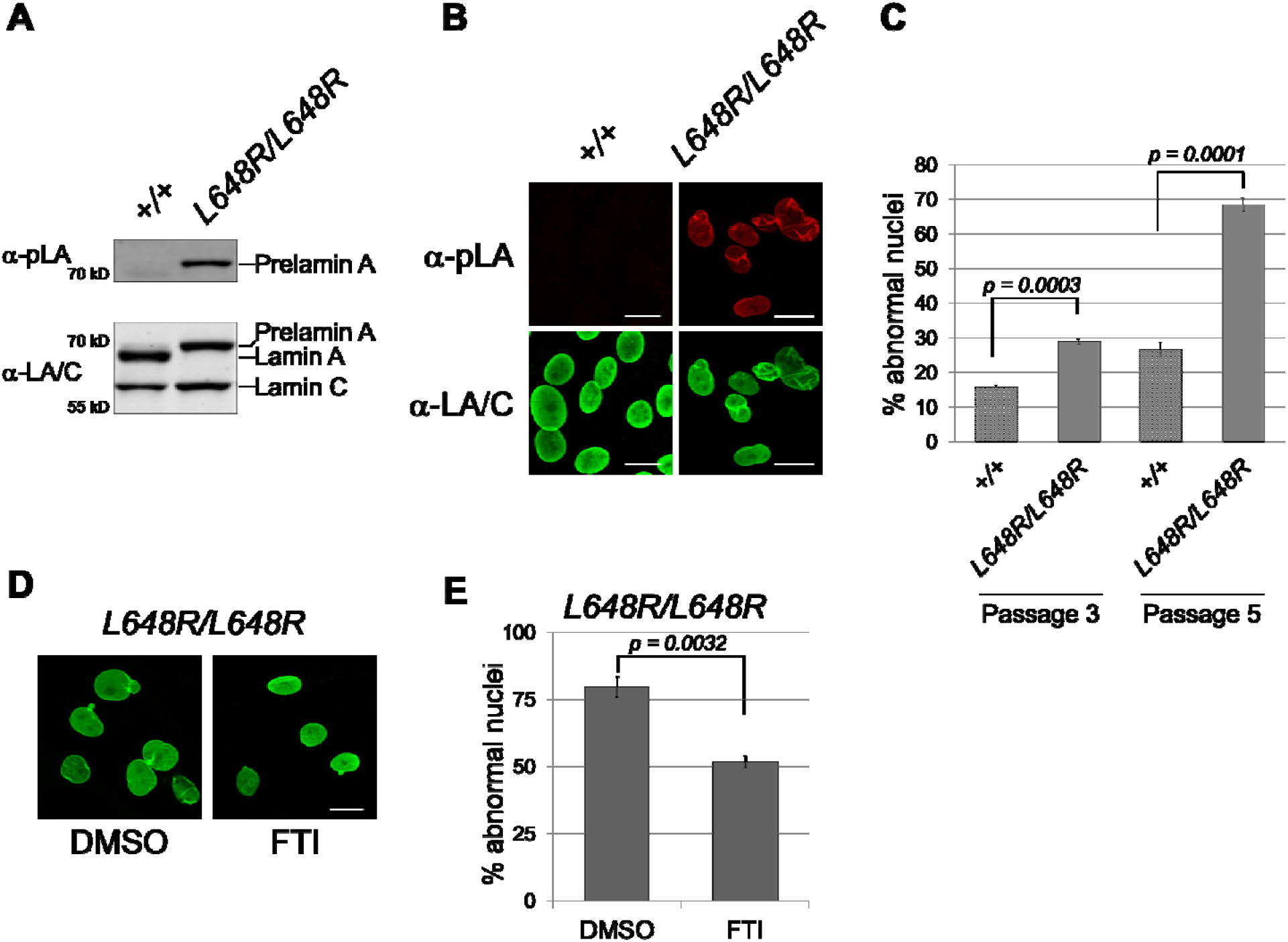
Farnesylation-dependent abnormal nuclear morphology in *Lmna*^L648R/L648R^ MEFs. (*A*) Immunoblots of protein extracts from MEFs generated from *Lmna*^+/+^ (*+/+*) and *Lmna*^L648R/L648R^ (*L648R/L648R*) mouse embryos. Blots were probed with an antibody specific for prelamin A (α-pLA, top) and anti-lamin A/C antibody that recognizes prelamin A, lamin A and lamin C (α-LA/C, bottom). (*B*) Immunofluorescence photomicrographs showing the indicated MEFs at passage 5 stained with antibodies that specifically recognize prelamin A (top) or prelamin A, lamin A and lamin C (bottom). Scale bar = 20 μm. (*C*) Quantification of aberrant nuclear morphology in MEFs at passage 3 and passage 6. Three independently grown MEF cultures were fixed, stained, and ~100 nuclei counted from each. Values are means (*N* = 3) and error bars indicate SEM. Representative images of nuclei that are counted as abnormal, including those that are blebbed, misshapen, or highly crenylated, are shown in *SI Appendix*, Fig. S5. (*D*) Immunofluorescence photomicrographs showing *Lmna*^L648R/L648R^ MEFs at passage 6 treated with dimethyl sulfoxidie (DMSO) vehicle or FTI as described in the Materials and Methods. Scale Bar = 20 μm. (*E*) Percentages of nuclei with aberrant nuclear morphology in *Lmna*^L648R/L648R^ MEFs treated with DMSO or FTI. For quantification, 3 independently grown MEF cultures were fixed, stained, and ~100 nuclei counted from each. Values are means (*N* = 3) and error bars indicate SEM. Statistical analyses were performed by unpaired, two-tailed *t*-test.

## Discussion

HGPS, MAD-B and other progeroid disorders arise from genetic mutations that lead to the accumulation of permanently farnesylated prelamin A or variant proteins (1–3). We have generated *Lmna*^L648R/L648R^ mice that accumulate a farnesylated prelamin A variant (and no mature lamin A) with a single amino acid substitution at the ZMPSTE24 cleavage site that blocks processing. Like mice with *Lmna* mutations that lead to production of progerin and *Zmpste24*^-/-^ mice that accumulate prelamin A, the *Lmna*^L648R/L648R^ mice develop progeroid phenotypes. Fibroblasts from these mice also have abnormal nuclear morphology that is reversible by pharmacological blockage of protein farnesylation, which is also the case for human and mouse cells expressing progerin or deficient in ZMPSTE24 (38). Notably, however, *Lmna*^L648R/L648R^ mice have dramatically greater longevity (~1.5 to > 2 years) as compared to *Zmpste24*^-/-^ mice (4-7 months) (9, 10) and *Lmna* mouse models of HGPS that have 2 mutant alleles encoding progerin (4-10 months) (26, 27, 29, 39). Because *Lmna*^L648R/L648R^ mice do not die prematurely, and can live more than 2 years, they are an ideal model for studying the effects of permanently farnesylated prelamin A in the context of physiologic aging.

The phenotype of *Lmna*^L648R/L648R^ mice in some ways resembles that of the one human patient described to date with the corresponding, albeit heterozygous, *LMNA* L647R mutation (25). The patient was small for chronological age, microcephalic, and had a “MAD-B like” condition with micrognathia, dental crowding, decreased subcutaneous fat and bone defects, including acro-osteolysis and other radiological findings suggestive of abnormal bone resorption or deposition. Fibroblasts from this patient expressed prelamin A to lamin A at a ratio of approximately 1:1; however, this individual was not nearly as severely affected as children with HGPS that are heterozygous for the *LMNA* G608G allele and whose fibroblasts generally express progerin to lamin A at a <1:1 ratio. Although the human data are from only one individual, they suggest that full-length prelamin A may be less “toxic” to cells than progerin, which contains a 50 amino acid deletion.

To a large extent, *Lmna*^L648R/L648R^ mice phenocopy what has been reported for *Zmpste24*^-/-^ mice in terms of growth, skull and other bone defects (9, 10, 12, 34). However, their phenotype is far less severe than that of *Zmpste24*^-/-^ mice and they do not exhibit premature lethality. Both mice have severe mandibular and dental abnormalities that could potentially lead to malnourishment causing the observed failure to thrive. However, *Lmna*^L648R/L648R^ mice have decreased body mass compared to wild type mice at ages before dental malocclusion becomes prominent, suggesting that factors other than dental abnormalities resulting in decreased caloric intake may contribute to their failure to thrive. *Lmna*^L648R/L648R^ mice also have decreased whole body fat content, as do *Zmpste24*^-/-^ mice and humans with MAD-B (7, 9, 10, 40, 41). Decreased body fat content of *Lmna*^L648R/L648R^ mice was not associated with insulin resistance. Female *Lmna*^L648R/L648R^ mice had modest hyperglycemia after glucose administration, but a lower than normal plasma insulin concentration. *Zmpste24*^-/-^ mice also have plasma insulin concentrations lower than wild type mice (42). Conditional deletion of *Zmpste24* from adipocytes leads to only modest body fat loss only in male mice (43). These findings suggest that decreased body fat in *Lmna*^L648R/L648R^ mice is not caused by a direct effect of prelamin A on adipocytes.

Why are *Lmna*^L648R/L648R^ mice similar to, but less severely affected than *Zmpste*24^-/-^ mice, given that both express solely farnesylated prelamin A and no mature lamin A? One possibility is that they are exposed to a lower “dose” of prelamin A. Z*mpste24*^-/-^ mice heterozygous for *Lmna* that express half the amount of prelamin A have significantly less severe phenotypes than those homozygous for *Lmna* (12, 13). However, *Lmna*^L648R/L648R^ mice like *Zmpste24*^-/-^ mice on a homozygous *Lmna* background express roughly equal amounts of prelamin A relative to lamin C, so this explanation seems unlikely (12, 13). Another possibility is that the single amino acid substitution renders the L648R variant less “toxic” than native prelamin A. This is difficult to reconcile with the wealth of data suggesting that it is the farnesyl moiety of prelamin A that is primarily responsible for its adverse effects in cultured cells and mice. Furthermore, mice engineered to express only a non-farnesylated prelamin A, without any lamin C, develop cardiomyopathy but do not have progeroid phenotypes and live much longer and have less severe growth retardation than *Zmpste24*^-/-^ mice (38). We favor the hypothesis that ZMPSTE24’s other crucial cellular functions, in addition to prelamin A cleavage, contribute to the severity of phenotypes and limited lifespan of the *Zmpste24*^-/-^ mice. ZMPSTE24 has a key role in clearing clogged translocons, as well as other less mechanistically well understood roles in viral defense, protein secretion, ER protein quality control, membrane stress, and establishing membrane protein topology (14–22). Loss of one or several of these other functions may make *Zmpste24*^-/-^ mice more susceptible to the “toxic” effects of prelamin A or create additional pathologies that exacerbate those resulting from prelamin A. *Lmna*^L648R/L648R^ mice also live significantly longer and have milder phenotypes than reported for various mouse models expressing progerin, the prelamin A variant in HGPS (26–29, 31, 37). Hence, the loss of 50 amino acids in progerin may make it more toxic than full-length prelamin A. In any case, much research has focused on *Zmpste24*^-/-^ and HGPS mouse models to understand the role of prelamin A in aging, the significant longevity of *Lmna*^L648R/L648R^ mice makes them an alternative, and potentially an ideal one to study how prelamin A impacts several aging tissues.

Brayson et al. (44) reported that transgenic cardiomyocyte-specific expression of human prelamin A with the L647R amino acid substitution causes cardiomyopathy in mice, with abnormal heart function detected as early as 4 weeks of age. In contrast, our mice expressing the mouse prelamin A L648R variant from the endogenous *Lmna* alleles lived almost 2 years without evidence of cardiomyopathy. Although only a few MAD-B patients have been thoroughly described in the literature (7), early-onset primary cardiomyopathy has not been described as a feature of these individuals. One patient with MAD-B and a compound heterozygous *ZMPSTE24* mutation reportedly has a greater carotid intima media thickness relative to chronological age, and a raised proteomic classifier for the diagnosis of coronary artery disease (45). Cardiac complications of cardiovascular disease, rather than primary cardiomyopathy, lead to early death in HGPS (46). Arteries from patients with HGPS show loss of medial smooth-muscle cells, adventitial thickening, and other pathological alterations that lead to sclerotic plaques in coronary arteries (47–50). This vascular pathology is the cause of myocardial infarctions and ischemic heart disease invariably present in individuals with HGPS. Several mouse models of HGPS develop vascular alterations that mimics this human pathology (26–30, 51). *Zmpste24*^-/-^ mice may die at too young an age to determine if prelamin A accumulation causes vascular pathology. *Lmna*^L648R/L648R^ mice therefore provide a novel model to determine the effects of prelamin A in the development of cardiovascular disease. We did not observe vascular pathology when we examined these mice at 7 months of age; however, we will continue to monitor them to see if it develops with advancing age or in combination with genetic susceptibility to atherosclerosis as has been done with HGPS model mice (51). Indeed, Ragnauth *et al*. (52) have reported that prelamin A accumulation is prevalent in the medial vascular smooth muscle cells of human arteries from old individuals, but not young ones, and in atherosclerotic lesions. This same group also reported that prelamin A was present in vascular smooth muscle cells in calcified arteries of subjects receiving renal dialysis, a model for vascular aging (53). This raises the possibility that prelamin A accumulation contributes to pathology associated with physiological aging. Because of the near-normal life-span of the *Lmna*^L648R/L648R^ mice, they will provide a robust model to determine the impact of prelamin A on the vasculature during aging.

## Materials and Methods

### Mice

The Institutional Animal Care and Use committee at Columbia University Irving Medical Center approved all protocols. We used recombination-mediated genetic engineering or recombineering to construct vectors for manipulation of the mouse genome (54). Bacterial artificial chromosome clone RP23-281P8 carrying *Lmna* was purchased from BACPAC Resource. We engineered the T>G transversion at *Lmna* c.2176 in exon 11, which changed codon 648 CTC encoding Leu to CGC encoding Arg. We then inserted a Frt-Neo-Frt (FNF) cassette into intron 11 (See also *SI Appendix*, Fig. S1*A*). A gene targeting vector was constructed by retrieving the 1.6-kb left homology arm (5’ to L648R mutation), the FNF cassette, and the 1.45-kb right homology arm (end of FNF cassette to 3’) into pMCS-DTA vector carrying the *Diphtheria* toxin alpha chain (DTA) negative selection marker. The FNF cassette conferred G418 resistance during gene targeting in KV1 (129B6 hybrid) embryonic stem (ES) cells and the DTA cassette provided an autonomous negative selection to reduce random integration events during gene targeting. Several targeted ES cell clones were identified and injected into C57BL/6J blastocysts to generate chimeric mice. Male chimeras were bred to C57BL/6J female mice to transmit the *Lmna*-L648R-FNF allele. The FNF cassette was then removed by crossing mice with the *Lmna*-L648R-FNF allele to B6.Cg-Tg(ACTFLPe)9205Dym/J mice (The Jackson Laboratory; Stock No: 005703). The offspring were crossed to wild type C57BL/6J mice and heterozygous offspring subsequently crossed to wild type C57BL/6J mice for another several genenrations before intercorssing male and female heterozygous mice to generate *Lmna*^L648R/L648R^, *Lmna*^+/L648R^, and *Lmna*^+/+^ mice used in the experiments on a >90% pure C57BL/6J background. Mice were housed in a barrier facility with 12/12 h light/dark cycles, fed a chow diet, and genotyped by PCR using genomic DNA isolated from tail clippings.

### Immunoblotting of proteins isolated from mouse tissues

Proteins were extracted from mouse tissues, separated by electrophoresis in SDS-polyacrylamide slab gels, transferred to nitrocellulose membranes, and analyzed by immunoblotting using methods described previously (55). Primary antibodies for immunoblotting were rabbit anti-lamin A/C (Santa Cruz) at 1:5,000 dilution, rat monoclonal anti-prelamin A 3C8 (56) at 1:3,000 dilution, and mouse anti-GAPDH (Ambion) at 1:3,000 dilution. Secondary antibodies were ECL-horseradish peroxidase-conjugated anti-rabbit, anti-rat and anti-mouse antibodies (GE Healthcare) used at a dilution of 1:5,000. Signals were detected using SuperSignal West Pico PLUS Chemiluminescent Substrate (Thermo Fisher Scientific) and Autoradiography film (LabScientific). For quantification, films were scanned with a digital scanner and the band signal intensities were quantified using Fiji ImageJ (https://imagej.net/Fiji) and analyzed using Excel (Microsoft).

### Growth and survival analyses

Mice were weighed twice per week in the first 12 weeks, once per week up to 40 weeks, and once every two weeks for remainder of their lifespans. For survival analyses, a combined endpoint of death or distress severe enough that a staff veterinarian blinded to genotype determined that euthanasia was necessary. Euthanasia was performed according to the approved protocol of the Institute of Comparative Medicine at Columbia University Irving Medical Center.

### Blood biochemical parameters and glucose tolerance tests

Mice were fasted for 5 hours, and blood was collected and centrifuged at 3,000g at 4 °C for 10 minutes. Plasma was collected and stored at −80 °C until analysis. Blood biochemistry analysis was performed on a Heska Element DC Chemistry Analyzer at the Institute of Comparative Medicine, Columbia University Irving Medical Center. Plasma insulin was measured using Ultra-Sensitive Mouse Insulin ELISA Kit (Crystal Chem). For glucose tolerance tests, mice were fasted in clean cages with no food and ad lib drinking water for 15 hours; then blood glucose was measured using One Touch Ultra (LifeScan) at time 0, and then 15, 30, 60, 90, and 120 minutes after intraperitoneal injection of a 2 g/kg bolus of sterile glucose solution.

### Micro-CT and image analysis

Mice and dissected bones were scanned using the Perkin-Elmer Quantum FX micro-CT imaging system (PerkinElmer) at the Oncology Precision Therapeutics and Imaging Core at Columbia University Irving Medical Center. For imaging mice, the animal was anesthetized with 2-3% isoflurane. The skeletal structure of the mouse was scanned in the axial plane placed in a cylindrical sample holder at the field of view of 20-60 mm for lower-high resolution. A single scan of 4 minutes was taken, and the animal then was removed from the micro-CT machine, returned to its cage, and kept warm for homoeothermic stability until consciousness returned. Dissected bone was placed on the sample holder and scanned at the same settings as for mice. Micro-CT VOX files were analyzed using Analyze 14.0 (AnalyzeDirect) and images were created using a surface rendering technique. The VOX files were also transferred to DICOM format and analyzed using AMIRA software. To obtain high-resolution images (6.6 μm), the dissected heads were scanned by SkyScan 1272 (Bruker BioSpin Corp). For mandible size analysis, the scanned 2D images were segmented to select the mandibular area, followed by rendering of a 3D model from the segmented images, and computed statistical information of the 3D model analyzed to obtain the structure measurements using AMIRA software (Thermo Fisher Scientific). For bone density analysis, the top 60 images of vertebra L5 were segmented by selecting the trabecular bone area in all 60 2D images, and then the 60 segmented images were used to construct a 3D surface model. The computed statistical information of the 3D model was used to obtain bone volume and non-bone volume, followed by statistical analysis.

### MEF isolation, culture, immunoblotting, immunofluorescence microscopy, and FTI treatment

Fibroblasts were isolated from 13.5-day embryos of female *Lmna*^+/L648R^ mice that were crossed with male *Lmna*^+/L648R^ mice using previously described methods (57). Primary MEFs were cultured in Dulbecco’s modified Eagle medium (Invitrogen or Thermo Fisher Scientific) containing 10% fetal bovine serum (FBS; Gemini) with 5% CO_2_ at 37°C and propagated in the same media containing penicillin-streptomycin-glutamine (1:100 dilution of Gibco Penicillin-Streptomycin-Glutamine 100X; Thermo Fisher Scientific).

For immunoblotting, SDS sample buffer was added to MEFs and the lysates sonicated and then heated for 10 min at 65°C. Proteins in lysates were separated by electrophoresis in 10% SDS-polyacrylamide slab gels and transferred to nitrocellulose using methods described previously (58), except that blocking was done with phosphate-buffered saline plus 0.1% Triton X-100 (PBS-T) containing 5% nonfat dry milk (Biorad Blotting grade Blocker) and 10 mM NaN_3_. Primary antibodies were mouse monoclonal anti-lamin A/C (Santa Cruz) at 1:1,000 dilution and rat monoclonal anti-prelamin A clone 3C8 (56) at 1:2,000 dilution. Secondary antibodies were goat anti-rat IRDye 680RD and goat anti-mouse IRDye 800C (LI-COR), and imaging was performed using a LI-COR Odyssey Imaging system.

For immunofluorescence microscopy, MEFs at the indicated passages were fixed in 4% freshly prepared formaldehyde for 20 min, permeabilized for 10 min in PBS-T, blocked for 1 hr in PBS-T containing 2% bovine serum albumin, and co-labeled with the mouse monoclonal anti-lamin A/C antibodies at 1:500 dilution and the rat monoclonal anti-prelamin A antibody 3C8 at 1:300 dilution for 1 hr. Cells were then incubated for 1 hr with IRDye680RD goat anti-mouse (LI-COR) and Alexa Fluor 647 donkey anti-rat (Abcam) secondary antibodies. Images were captured on a ZEISS LSM700 system with ZEISS Plan-NEOFLUAR 40x/1.3 oil submersion objective and analyzed by ZEN (black edition) software. Florescence intensities were quantified using Fiji ImageJ and analyzed using Excel. To test the effects of a FTI, MEFs were treated for 48 hr with 2 μg/ml lonafarnib (The Progeria Research Foundation) dissolved in DMSO, or DMSO only as the vehicle control.

### Statistics for mouse experiments

For comparisons among more than two groups, ANOVA and Tukey’s post hoc multiple comparisons were carried out using GraphPad Prism 5. Student’s *t*-tests of comparisons between two means were performed using Excel 2016 (Microsoft); bar graphs and body mass growth curves were generated using the same software. The Kaplan-Meier estimator in statistical software GraphPad Prism 5 (Prism Software) was used to generate the survival curves.

## ACKNOWLEDGEMENTS

We thank Drs. Loren Fong and Stephen G. Young (University of California, Los Angeles) for antibodies against prelamin A and the Genetically Modified Mouse Model Shared Resource of the Herbert Irving Comprehensive Cancer Center at Columbia University for assistance in generating mice. Research reported in this publication was supported by the National Institute of Aging and National Institute of Dental and Craniofacial Research of the National Institutes of Health under award numbers R21AG058032 to S.M. and H.J.W, and R01DE015654 and DE026936 to W.H. The content is solely the responsibility of the authors and does not necessarily represent the official views of the National Institutes of Health.

