## Supplemental Materials for "Abolishing the prelamin A ZMPSTE24 cleavage site leads to progeroid phenotypes with near-normal longevity in mice"

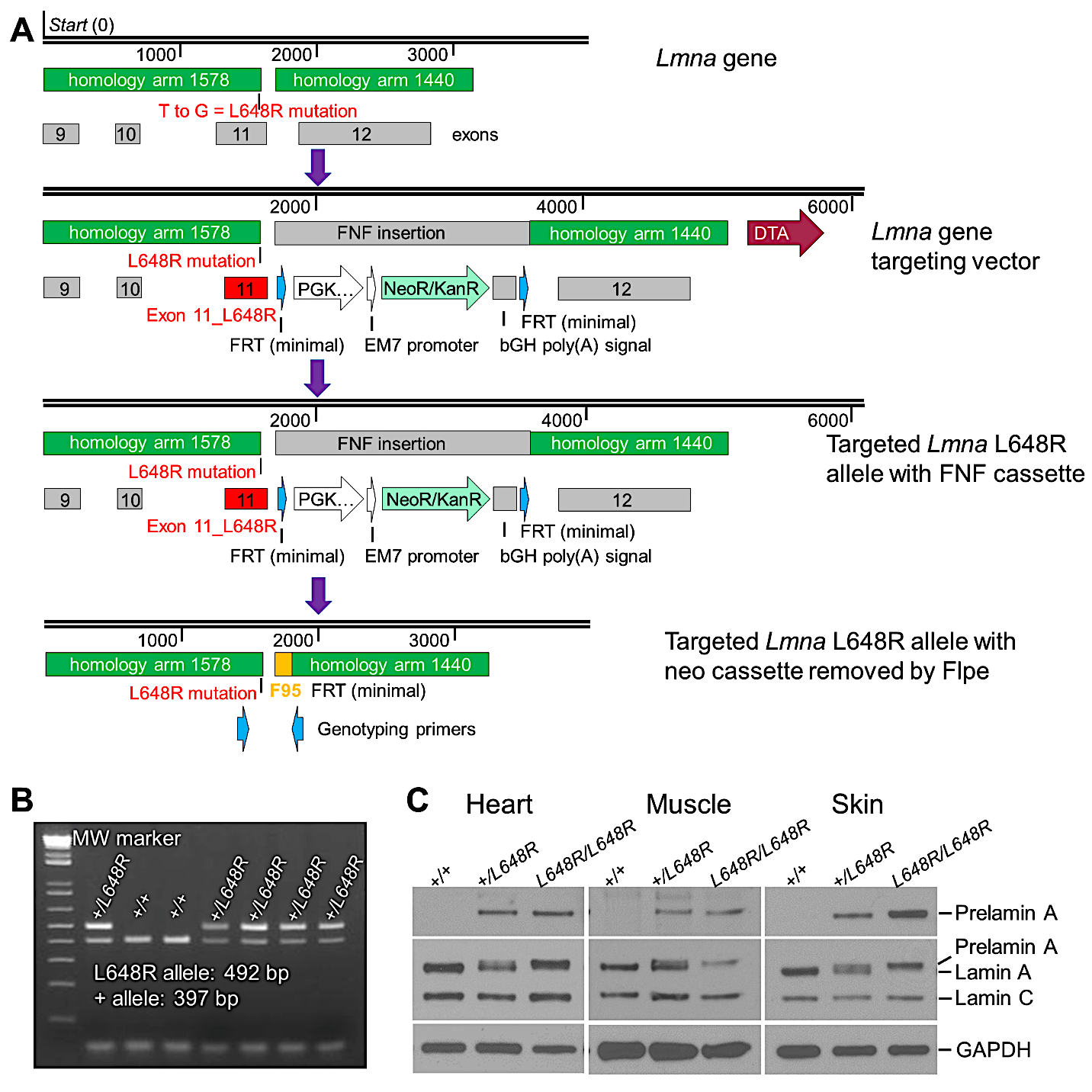

Fig. S1. Generation of mice with a *Lmna* L648R allele. (*A*) Schematic diagram showing procedure used to generate the *Lmna* L648R allele by recombineering. See Materials and Methods above for additional details. (*B*) Ethidium bromide-stained agarose gel showing PCR-amplified DNA from *Lmna*^+/648R^ (*+/L648R*) founder mice and littermate *Lmna*^+/+^ (*+/+*) mice. The amplified fragment of the *Lmna* L648R allele is 492 base pairs (bp) and the wild type (*+*) allele 397 bp. This is because the mutant allele contains the F95 DNA fragment shown in *A*. The left lane of the gel (MW marker) shows an Invitrogen 1 Kb Plus DNA Ladder. (*C*) Immunoblots of protein extracts from heart, skeletal muscle (Muscle) and skin of *+/+*, *+/L648R* and *Lmna*^L648R/L648R^ (*L648R/L648R*) mice. Blots were probed with an antibody specific for prelamin A (top), an anti-lamin A/C antibody that recognized prelamin A, lamin A and lamin C (middle) or anti-GAPDH antibody as loading control (bottom).

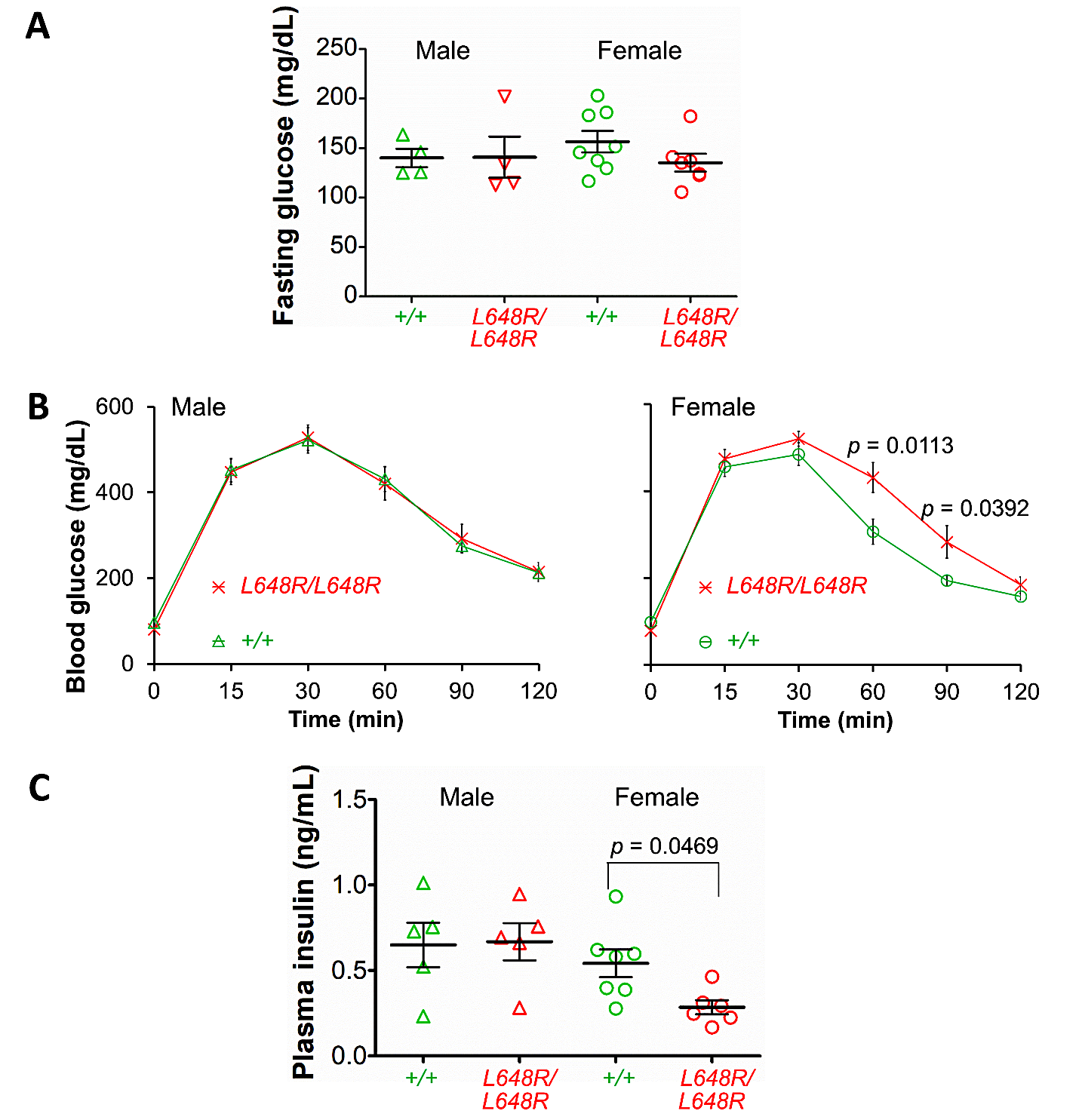

**Fig. S2**. Blood glucose and insulin concentrations in *Lmna*^L648R/L648R^ mice. (*A*) Fasting blood glucose concentrations in male and female *Lmna*^+/+^ (*+/+*) and *Lmna*^L648R/L648R^ (*L648R/L648R*) mice at 30 weeks of age. Each triangle or circle represents value for an individual animal; long horizontal bars represent mean and errors bars indicate SEM. (*B*) Blood glucose concentration versus time before and after injection of a glucose bolus in overnight-fasted *L648R/L648R* and *+/+* mice at 35 weeks. Values are means and error bars indicate SEM (*N* = 8 per genotype for male mice, and *N* = 12 per genotype for female mice). (*C*) Fasting plasma insulin concentrations in 30-week-old male and female *+/+* and *L648R/L648R* mice. Each triangle or circle represents value for an individual mouse; long horizontal bars represent means and errors bars indicate SEM.

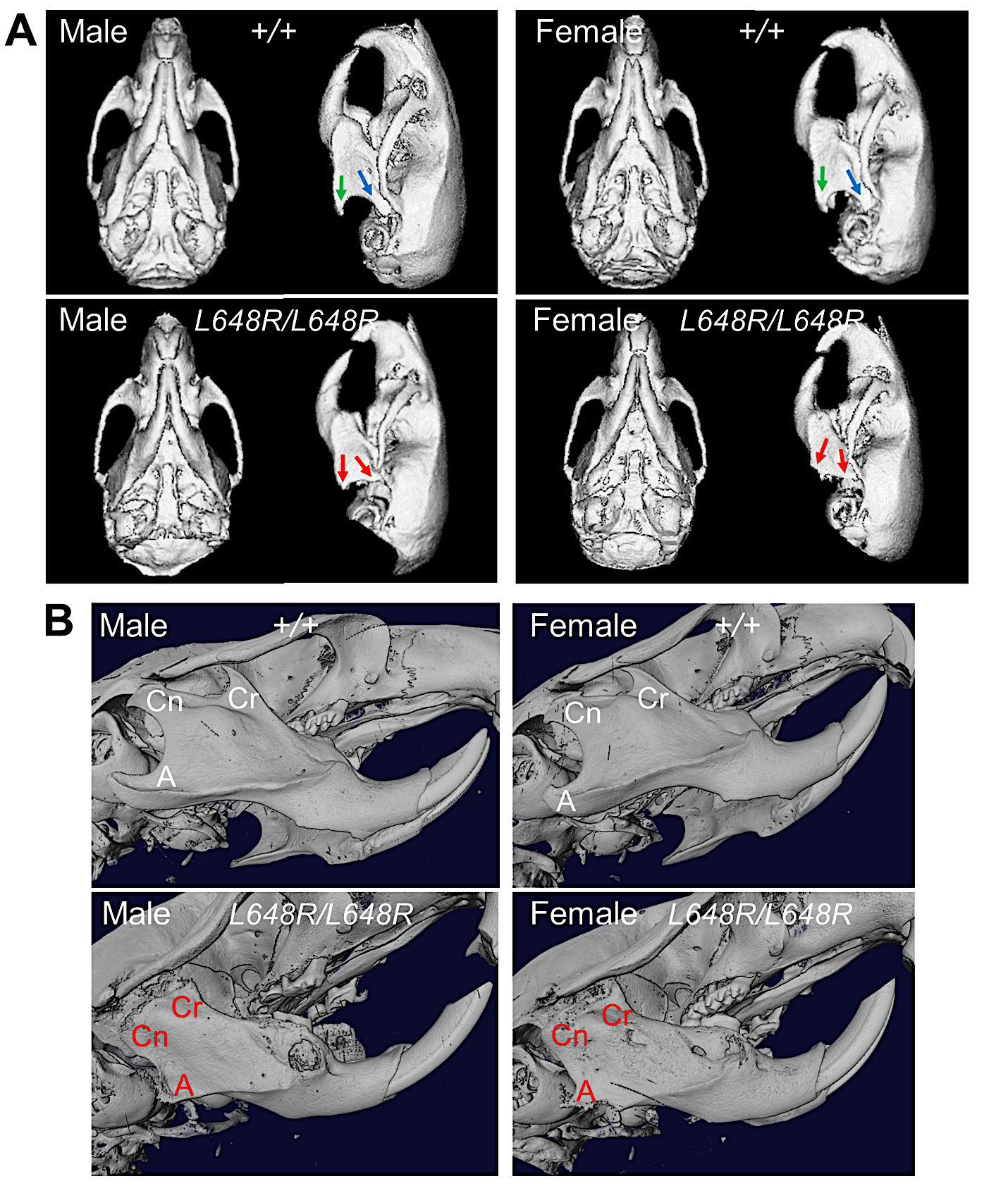

**Fig. S3**. Micro-CT images of skulls of *Lmna*^L648R/L648R^ mice at approximately 30 weeks of age. (*A*) Micro-CT-generated images showing representative ventral and left lateral views of skulls of living male and female *Lmna*^+/+^ (*+/+*) and *Lmna*^L64R/L648R^ (*L648R/L648R*) mice. In *+/+* mice, green arrows indicate mandibular angular process and blue arrows condylar process. These processes are smaller in *L648R/L648R* mice (red arrows). (*B*) Micro-CT-scanned and 3D-reconstructed images of skulls from male and female *+/+* and *L648R/L648R* mice. Cr: coronoid process; Cn: condylar process; A: angular process. Red lettering indicates degeneration of these processes in the mutant mice.

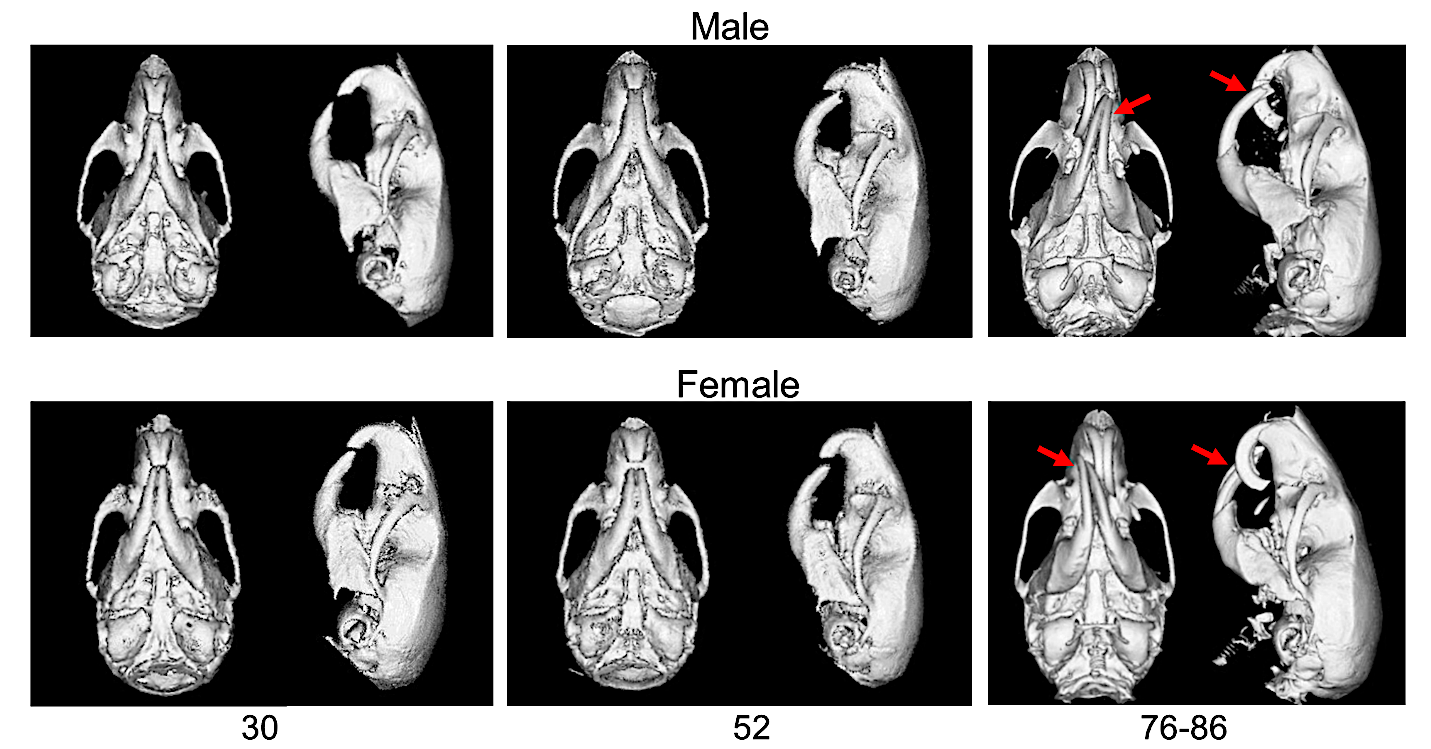

**Fig. S4.** Radiological confirmation of dental malocclusion in older *Lmna*^L648R/L648R^ mice. Micro-CT-scanned and 3D-rendered images showing representative ventral and left lateral views of skulls of living male and female *Lmna*^L648R/L648R^ mice at the ages indicated in weeks confirming dental malocclusion (red arrows).

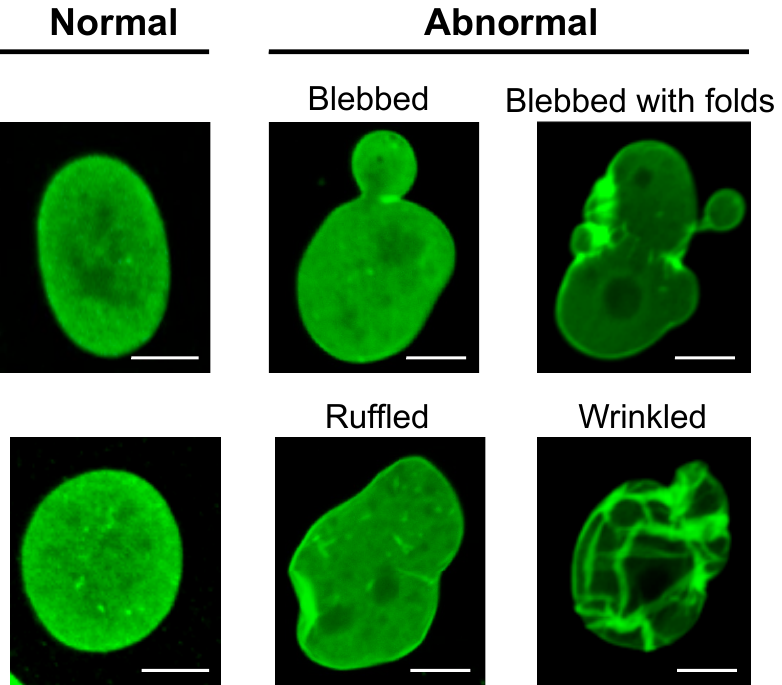

**Fig. S5.** Gallery of representative normal and abnormal nuclear morphologies observed in *Lmna*^L648R/L648R^ MEFs. Immunofluorescence photomicrographs of *Lmna*^L648R/L648R^ MEFs labeled with anti-lamin A/C antibodies show morphologically normal ovoid or round nuclei with smooth edges (left) and abnormal nuclei with a variety of indicated shapes and characteristics (right). These criteria were used to determine the percentage of abnormal nuclei in Fig. 5. Scale bars: 10 µm.

**Table S1.** Blood biochemical parameters in 30-week-old male and female *Lmna*^+/+^ and *Lmna*^L648R/L648R^ mice.

|  |  | Male | | Female | |
| --- | --- | --- | --- | --- | --- |
| Parameter | Reference Ranges | *Lmna*^+/+^ | *Lmna*^L648R/L648R^ | *Lmna*^+/+^ | *Lmna*^L648R/L648R^ |
| Total protein | 3.5-7.2 g/dL | 5.64 ± 0.12 | 5.52 ± 0.09 | 5.47 ± 0.09 | 5.60 ± 0.13 |
| Albumin | 2.5-3.4 g/dL | 2.36 ± 0.07 | 2.50 ± 0.08 | 2.37 ± 0.10 | 2.57 ± 0.06 |
| Alkaline phosphatase | 35-96 U/L | 79.80 ± 5.43 | 79.40 ± 3.68 | 118.00 ± 12.72 | 136.33 ± 16.88 |
| Total bilirubin | 0-0.9mg/dL | 0.30 ± 0.08 | 0.15 ± 0.04 | 0.12 ± 0.02 | 0.10 ± 0 |
| Phosphorus | 5.7-9.2 mg/dL | 10.48 ± 1.38 | 9.40 ± 0.97 | 8.69 ± 0.43 | 7.17 ± 0.82 |
| Cholesterol | 40-130 mg/dL | 94.00 ± 8.00 | 77.00 ± 3.61 | 102.20 ± 8.07 | 74.67 ± 6.74 |
| Alanine aminotransferase | 17-77 U/L | 46.80 ± 3.65 | 56.00 ± 16.93 | 39.43 ± 4.71 | 36.33 ± 3.98 |
| Calcium | 7.1-10.1 mg/dL | 8.24 ± 0.48 | 8.42 ± 0.50 | 8.30 ± 0.38 | 7.95 ± 0.38 |
| Creatinine | 0.2-0.9 mg/dL | 0.36 ± 0.07 | 0.35 ± 0.09 | 0.34 ± 0.03 | 0.40 ± 0.07 |
| Blood urea nitrogen | 8-33mg/dL | 22.76 ± 0.64 | 22.28 ± 1.01 | 23.34 ± 2.01 | 24.45 ± 1.10 |
| Triglyceride | 0-0 mg/dL | 78.00 ± 5.45 | 71.40 ± 5.88 | 70.14 ± 3.06 | 71.83 ± 5.53 |

Values are means ± SEM. *N* = 5 for male *Lmna*^+/+^ and *N* = 5 for male *Lmna*^L648R/L658R^ mice; *N* = 7 for female *Lmna*^+/+^ and *N* = 6 for female *Lmna*^L648R/L658R^ mice.
